# Nse5/6 is a negative regulator of the ATPase activity of the Smc5/6 complex

**DOI:** 10.1101/2021.02.12.430902

**Authors:** Stephen T. Hallett, Pascale Schellenberger, Lihong Zhou, Fabienne Beuron, Ed Morris, Johanne M. Murray, Antony W. Oliver

## Abstract

The multi-component Smc5/6 complex plays a critical role in the resolution of recombination intermediates formed during mitosis and meiosis, and in the cellular response to replication stress. Using recombinant proteins, we have reconstituted a series of defined *S. cerevisiae* SMC5/6 complexes, visualised them by negative stain electron microscopy, and tested their ability to function as an ATPase. We find that only the six protein ‘holo-complex’ is capable of turning over ATP and that its activity is significantly increased by the addition of double-stranded DNA to reaction mixes. Furthermore, stimulation is wholly dependent on functional ATP-binding pockets in both Smc5 and Smc6. Importantly, we demonstrate that budding yeast Nse5/6 acts as a negative regulator of Smc5/6 ATPase activity, binding to the head-end of the complex to suppress turnover, irrespective of the DNA-bound status of the complex.

## INTRODUCTION

The central scaffold of each eukaryotic Structural Maintenance of Chromosomes (SMC) complex is formed by an obligate heterodimer, via specific parings of the Smc1 + Smc3, Smc2 + Smc4, and Smc5 + Smc6 proteins, creating cohesin, condensin and Smc5/6 respectively. Globular domains found at the N- and C-termini of each SMC protein, contain Walker A and Walker B ATP-binding motifs that are brought together in space to form a so-called ‘head’ domain (**see Figure 1**). The two halves of the resulting ATPase are connected via a long anti-parallel coiled coil insertion (‘arm’) that is capped by a ‘hinge’ domain; the hinge serving to both reverse the directionality of the coiled coil as well as provide the major dimerisation interface between the two SMC proteins^1–4^. A second, more transitory interface is created between the head domains of the two SMC proteins, regulated through binding and hydrolysis of ATP [for recent comprehensive reviews, please see refs. 5,6].

**Figure 1.**
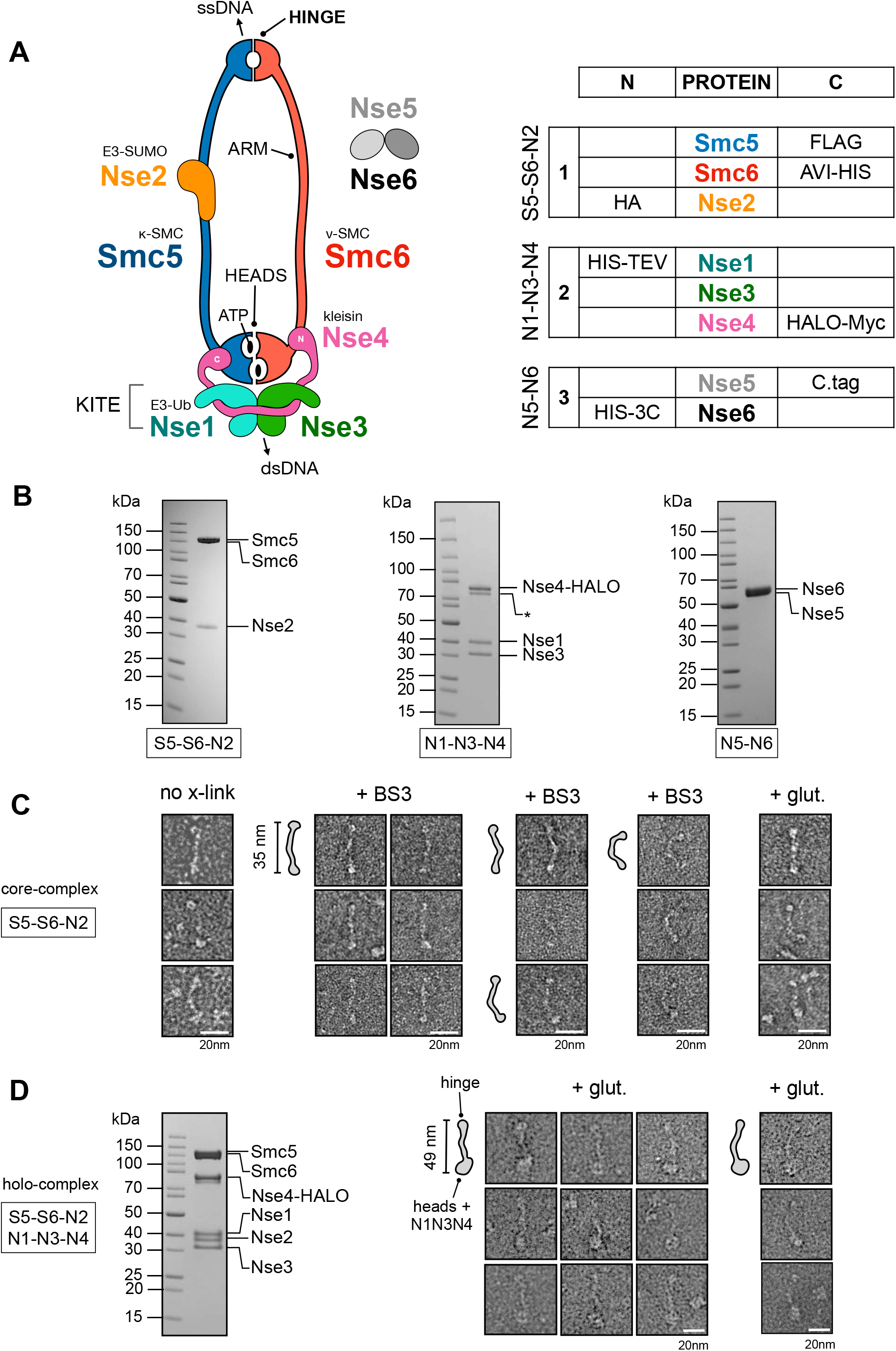
Purification and visualisation of the *S. cerevisiae* Smc5/6 complex. (**A, left**) Cartoon schematic showing the known subunits of the *Saccharomyces cerevisiae* Smc5/6 complex and their spatial relationships. A heterodimer of Smc5 and Smc6 is formed through an obligate interface formed at the so-called ‘hinge’. Nse2, a SUMO E3-ligase binds to the coiled coil ‘arm’ of Smc5. The kleisin subunit Nse4, serves to connect and join the Nse1 / Nse3 KITE subcomplex (kleisin-interacting tandem winged-helix elements) to the ATP-binding ‘heads’ of Smc5 and Smc6, through separate interactions at its N- and C-termini; providing the κ- and ν-SMC designation for Smc5 and Smc6, respectively. The Nse1 subunit provides Ubiquitin E3-ligase activity. Two additional subunits, Nse5 and Nse6 form an obligate heterodimer that can bind to the Smc5/6 holo-complex. Single-stranded DNA-binding activity has been shown for the hinge region of Smc5/6 ^1^, whilst a separate and distinct double-stranded DNA binding activity has been shown for the Nse3 subunit^58^ (**A, right**) Details of components in each baculovirus construct generated using the biGBac system^59^: S5-S6-N2 (1; pBIG1a), N1-N3-N4 (2; pBIG1b) and N5-N6 (3; pBIG1c) and their respective N or C-terminal affinity / epitope tags. (**B**) Representative colloidal-blue stained SDS-PAGE gels for each of the indicated subcomplexes. (**C**) Representative images of particles, from micrographs of uranyl acetate negative-stained S5-S6-N2, with either no crosslinking (left) or mild-crosslinking with BS3 or glutaldehyde (glut.). (**D**) Colloidal-blue stained SDS-PAGE gel of the purified Smc5/6 holo-complex (S5-S6-N2 / N1-N3-N3; pBIG2ab). (**E**) Representative images of particles, taken from micrographs of uranyl acetate negative-stained holo-complex. Particle outlines are provided to aid visualisation, with overall lengths estimated from micrographs for extended ‘I’-conformations of S5-S6-N2 and holo-complex.

Each SMC complex is then elaborated by binding of additional protein subunits highly specific to each family member. In the Smc5/6 complex, these subunits are designated as Non-SMC-Elements (NSMCE in humans, Nse in yeasts^7^). The SMC5/6 ‘holo-complex’ includes four such subunits: Nse2, an E3 SUMO Ligase, which associates with the arm of Smc5; plus Nse1 (E3 Ubiquitin Ligase), Nse3 (MAGE) and Nse4 (kleisin) that coalesce to form a defined sub-complex that binds to the head domains (**Figure 1**).

The precise cellular roles of the Smc5/6 complex remain enigmatic, but mutations within its component proteins have clear and definite impacts on DNA replication and DNA damage repair processes, in particular acting to suppress inappropriate homologous recombination structures that can be formed when replication forks stall or collapse, as well as assisting in resolution of sister chromatids during meiosis [reviewed in refs. 8-11]. Smc5/6 is also known to be a viral restriction factor for hepatitis B^12,13^, herpes simplex-1^14^ and papillomavirus type 31 viruses^15,16^. Furthermore, mutations found within the coding sequences of Nse2 and Nse3 have also been linked to human disease^17,18^.

Two additional protein subunits are also known to associate with the Smc5/6 holo-complex: Nse5 and Nse6 form an obligate heterodimer (Nse5/6) and both proteins are essential for viability in a range of different organisms, with the notable exception of fission yeast^19,20^. Identification of Nse5/6 orthologues has, however, been complicated by the lack of amino acid sequence identity in both proteins across different species. There is also no unifying or clear consensus with respect to their predicted domain composition, although structure-prediction programs have indicated that in the yeasts Nse6 may contain alpha-helical solenoids belonging to either the armadillo or HEAT repeat family^20^. The ‘functional equivalent’ of Nse5/6 in humans, formed by the SLF1/SLF2 heterodimer (Smc5/6 localisation factors 1 and 2) has been identified in a proteomics-based approach examining proteins recruited to psoralen-crosslinked chromatin^21^.

Nse5/6 is thought to promote recruitment to, or ‘loading’ of, the Smc5/6 complex onto chromatin^22,23^ in a manner similar to that described for the cohesin ‘loader complex’ Scc2-Scc4^20^ [reviewed in refs: 6,24]. In support of this hypothesis: *S. pombe* cells lacking Nse5/6 display a drastic reduction in the amount of Smc5/6 associated with chromatin^22,25^; in *S. cerevisiae*, hypomorphic mutations of Nse5 lead to reduced levels of Smc5/6 associated with stalled replication forks^26^; in humans the SLF1-SLF2 complex has been shown to recruit Smc5/6 to collapsed replication forks^21^.

At least part of the Smc5/6-recruitment function appears to be mediated by interaction of Nse5/6 with the multi-BRCT scaffold protein Rtt107 (Brc1 in *S. pombe*)^27^, which itself is recruited to sites of DNA damage via binding of its C-terminal BRCT-pair to γH2A. Notably, in fission yeast, Brc1 is a dosage compensator for a hypomorphic mutation of Smc6 known as *smc6-74* that leads to sensitivity to genotoxic agents and reduced levels of chromatin loading of Smc5/6^25,28^. Suppression is however dependent on the activity of the structure specific endonucleases Slx1-Slx4 and Mus81-Eme1^29^. A recent X-ray crystal structure of the N-terminal region of Rtt107 has revealed it to contain an unusual tetra-BRCT arrangement that can bind to a recognition sequence found at the N-terminus of Nse6, as well as motifs found in Mms22 and Slx4^30^. Thus, overexpression of Brc1 likely leads to increased recruitment of either Smc5/6 and/or the aforementioned structure-specific nucleases. However, it still remains unclear as to how binding of the Nse5/6 heterodimer to Smc5/6 promotes its ‘loading’ and retention on chromatin. There is also some ambiguity as to where its binds, with data from different organisms and laboratories supporting binding of Nse5/6 to the hinge, arms, or head regions of Smc5/6^31–33^.

Here we demonstrate that only the six protein Smc5/6 ‘holo-complex’ is capable of turning over ATP and that this activity is preferentially stimulated in the presence of dsDNA. We identify Nse5/6 as a negative regulator of the ATPase activity of the *S. cerevisiae* Smc5/6 complex, that is insensitive to the DNA-bound status of the complex. Furthermore, association of Nse5/6 induces a conformational change consistent with dis-engagement of the head-domains to prevent ATP-hydrolysis, compatible with its binding to the head-end of the Smc5/6 complex.

## RESULTS

### Reconstitution of the S. cerevisiae Smc5/6 complex

We co-expressed each of the known subunits of the *Saccharomyces cerevisiae* Smc5/6 complex in insect cells, initially as three distinct sub-complexes: (1) Smc5, Smc6, Nse2 (S5-S6-N2); (2) Nse1, Nse3, Nse4 (N1-N3-N4); and (3) Nse5, Nse6 (N5-N6) (**Figure 1B**). Using standard chromatographic techniques, we purified each sub-complex and visualised them on colloidal blue-stained SDS-PAGE gels **(Materials and Methods**, **Figure 1B**). We confirmed the identity of each protein component and its migration position on SDS-PAGE gels, by western blot using the various affinity / epitope tags incorporated into their respective expression cassettes (**Supplementary Figure 1**). Nse3, which is untagged, migrates at its expected size of 34 kDa (**Figure 1C, middle**).

### Negative stain transmission electron microscopy

The S5-S6-N2 complex proved to be relatively unstable (under the experimental conditions tested) but mild cross-linking with either glutaldehyde or BS3 greatly improved stability and aided visualisation of the complex by uranyl acetate negative stain electron microscopy (**Materials and Methods, Figure 1C**). The majority of S5-S6-N2 particles adopt an ‘arms-together’, ‘rod-like’ or ‘I’-conformation that has been seen in other exemplars of the SMC-family, i.e., an extended, predominantly linear conformation with an overall length of ~35 nm (**Figure 1C, middle**). However, some level of flexibility is still evident, as indicated by the appearance of the selected particles presented in the right-hand panel of **Figure 1C**. To produce the intact Smc5/6 ‘holo-complex’ we combined sub-complexes 1 and 2 into a single baculovirus (S5-S6-N2/N1-N3-N4). Again, we were readily able to purify, cross-link and visualise the resultant complex by negative stain electron microscopy (**Materials and Methods, Figure 1D**).

### 2D class averages and 3D model

Manual picking yielded 5397 particles with good levels of staining. Iterative classification and particle selection through successive 2D-averages, yielded 4301 particles that were subsequently used to generate a 3D model (**Materials and Methods, Table 1**). The resultant 2D class averages and 3D model (**Figure 2**) had sufficient features to allow identification of the hinge and head ends, and to determine an overall length of ~48 nm for the holo-complex. A protruding lobe of density, found approximately halfway along the length of the complex, is compatible with both the volume and expected position of Nse2 bound to the arm of Smc5 (**Figure 2 inset**; PDB: 3HTK^34,35^).

**Table 1.** Data summary for Transmission Electron Microscopy.

|  | **Holo-complex** | **Super-complex** | **MBP-labelled**<br>**Super-complex** |
| --- | --- | --- | --- |
| Figure # | 2 | 7E | 7H, S4 |
| Components | S5-S6-N2<br>N1-N3-N4 | S5-S6-N2<br>N1-N3-N4<br>+ N5-N6 | S5-S6-N2<br>N1-N3-N4<br>+ N5-N6 (MBP) |
| Microscope | Tecnai F20 | JEOL JEM-1400-plus |  |
| Detector | TemCam F416<br>CMOS (TVIPS) | OneView Camera (Gatan) |  |
| Voltage (kV) | 200 | 120 |  |
| Magnification | 50,000 | 60,000 |  |
| Pixel size (Å/pix) | 1.732 | 1.8 |  |
| Grid type | Quantifoil R1.2/1.3<br>copper 300 mesh<br>grids (Agar Scientific)<br>with an additional<br>layer of supporting<br>carbon added in<br>house | Continuous Carbon Film on 300 mesh<br>copper grids (Agar Scientific) |  |
| # Micrographs | 416 | 234 | 102 |
| # Particles picked | 5397 | 4469 | 5279 |
| # Particles in 3D model | 4301 | 3107 | 3990 |
| nominal resolution (Å); as reported by RELION / 3D-refine |  |  |  |
| Overall | 29 | 33 | 31 |
| “Hinge” | 25 | 28 | 30 |
| “Arm” | 22 | 31 | 27 |
| “Heads” | 27 | 28 | 31 |

**Figure 2.**
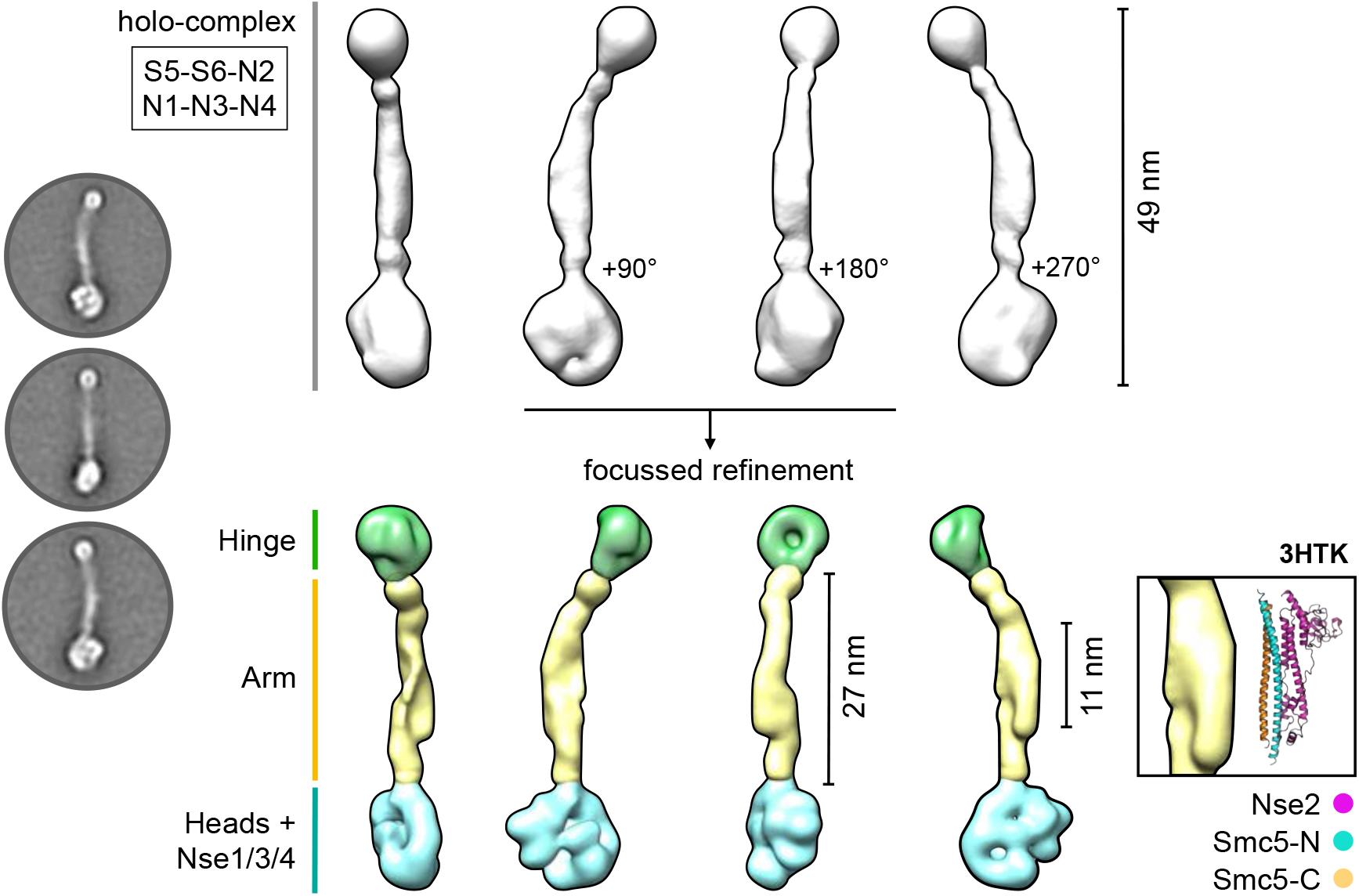
2D class averages and 3D model of the Smc5/6 holo-complex. (**Left**) Representative 2D class averages (**Top right**) Initial 3D reconstruction. (**Bottom Right**) 3D reconstruction from focussed refinement; splitting the model into masked sections corresponding to the ‘hinge’, ‘arm’ and ‘heads + NSE1/3/4’ regions of the holo-complex (coloured in green, yellow and cyan respectively). (**Inset**) A section of density protruding from the ‘arm’ region is consistent with the dimensions of the X-ray crystal structure of budding yeast Nse2 in complex with a short section of the Smc5 coiled-coil ‘arm’ (PDB ID: 3HTK). See associated key for additional details.

However, we noted that detail evident in the 2D class averages was not fully represented by the final model. To improve this, we pursued a focussed refinement strategy, splitting the holo-complex into three separate sections corresponding to the Hinge, Arm, and Heads + NSE1/3/4 to produce a segmented 3D model that better recapitulated the detail evident in 2D classes (**Figure 2 bottom**).

### ATPase activity assays with defined complexes

We first validated a NADH-coupled regenerative ATPase assay^36^ using recombinant RecQ5-HD (helicase domain) as a positive control, observing robust stimulation of ATP turnover by addition of a single-stranded 48mer oligonucleotide to the reaction mix (**Materials and Methods**, **Figure 3A**). We then examined the ATPase activity of purified S5-S6-N2 and S5-S6-N2/N1-N3-N4 complexes.

**Figure 3.**
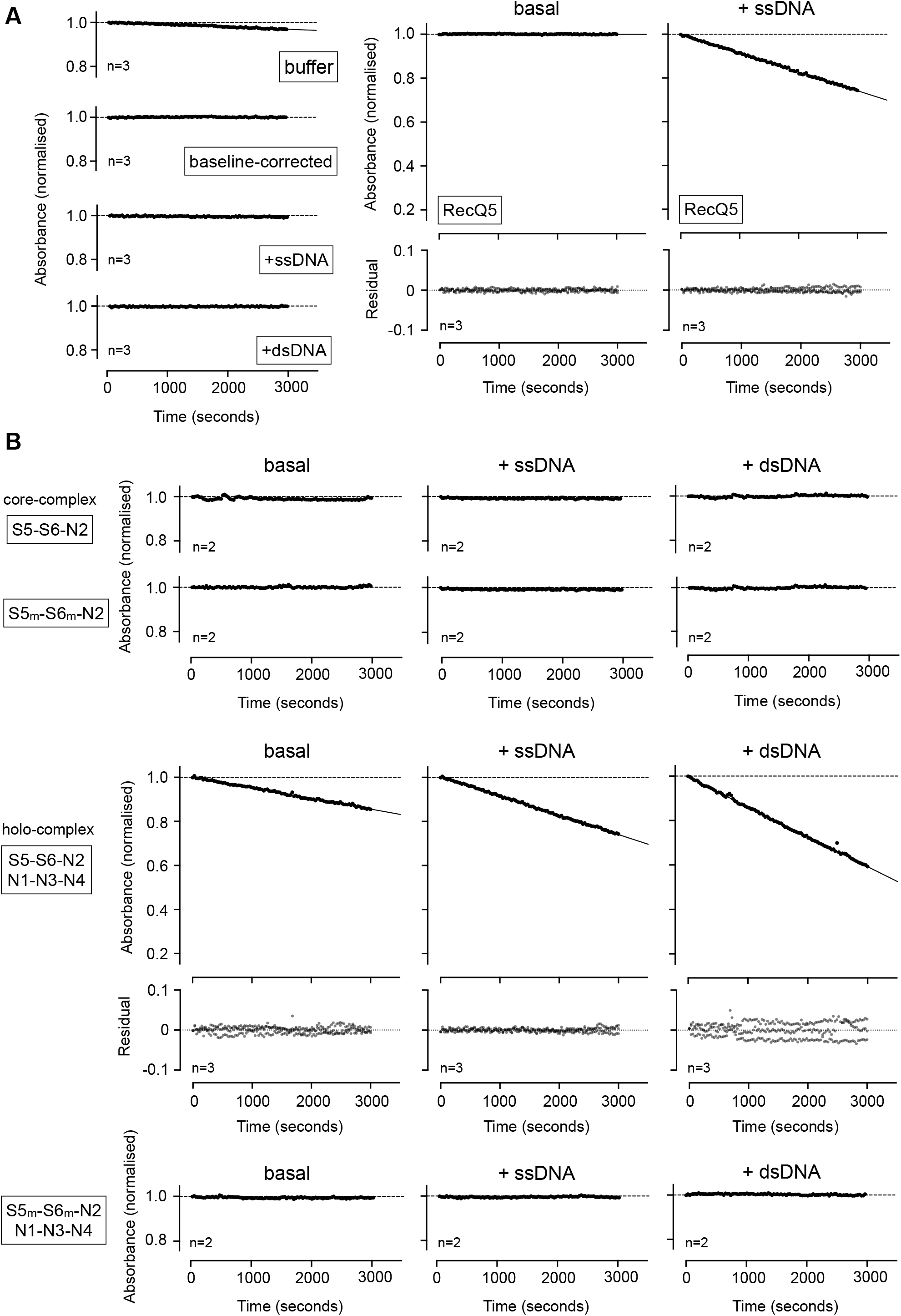
Measuring ATP-hydrolysis by defined Smc5/6 complexes. (**A, left**) Control experiments for the NADH-coupled regenerative ATPase assay. A highly consistent and reproducible baseline-drift is observed with buffer only controls [buffer], which can be corrected for by simple subtraction [baseline-corrected]. All subsequent data have been baseline-corrected in this manner. Addition of nucleic acid to the experimental buffer does not affect the baseline level of ATP consumption. (**A, right**) Addition of a single-stranded DNA oligonucleotide (48-mer) stimulates ATP turnover by recombinant human RecQ5 (helicase domain). (**B**) Neither purified S5-S6-N2, nor S5m-6m-N2 (containing inactivating Walker B mutations in both SMC subunits; respectively) turn over ATP. No stimulation of activity is observed with the addition of nucleic acid. (**C**) Purified Smc5/6 holocomplex (S5-S6-N2 / N1-N3-N4) has a basal level of ATP-turnover that can stimulated by the addition of nucleic acid. The equivalent inactivated complex (S5m-S6m-N2 / N1-N3-N4) cannot turn over ATP. In each case, plotted data represent the mean of the indicated number of experiments (n=X). For experiments showing activity, residual plots are also shown, generated by fitting experimental data to a straight-line equation by non-linear regression (solid black line); residuals are shown for data points generated in each experimental repeat (filled circles, coloured light, medium and dark grey). 5_m_ = Smc5-E1015Q, 6_m_ = Smc6-E1048Q.

Whilst we saw hydrolysis by the holo-complex, S5-S6-N2 was inactive, even when tested at a 7-fold higher concentration (**Figure 3B, Supplementary Figure 2**). Furthermore, whilst addition of ssDNA or dsDNA to the reaction mix clearly increased turnover by the holo-complex it did not elicit any effect on S5-S6-N2. Moreover, as hydrolysis could not be detected in purified complexes containing Walker B catalytic site mutations in both SMC subunits (5_m_ = E1015Q, 6_m_ = E1048Q) we were confident that the observed activity was not attributable to any potential co-purifying contaminants. Titration experiments then allowed quantification of the effects of adding either dsDNA or ssDNA to the holo-complex (**Figure 4A**). For titrations with dsDNA, the resultant activity curve approached a maximum plateau, with a calculated EC_50_ value of ~1.7 μM (**Figure 4B**). Over the concentration range tested, addition of ssDNA was far less stimulatory and a value for EC_50_ could only be estimated at (or above) a value of 16 μM.

**Figure 4.**
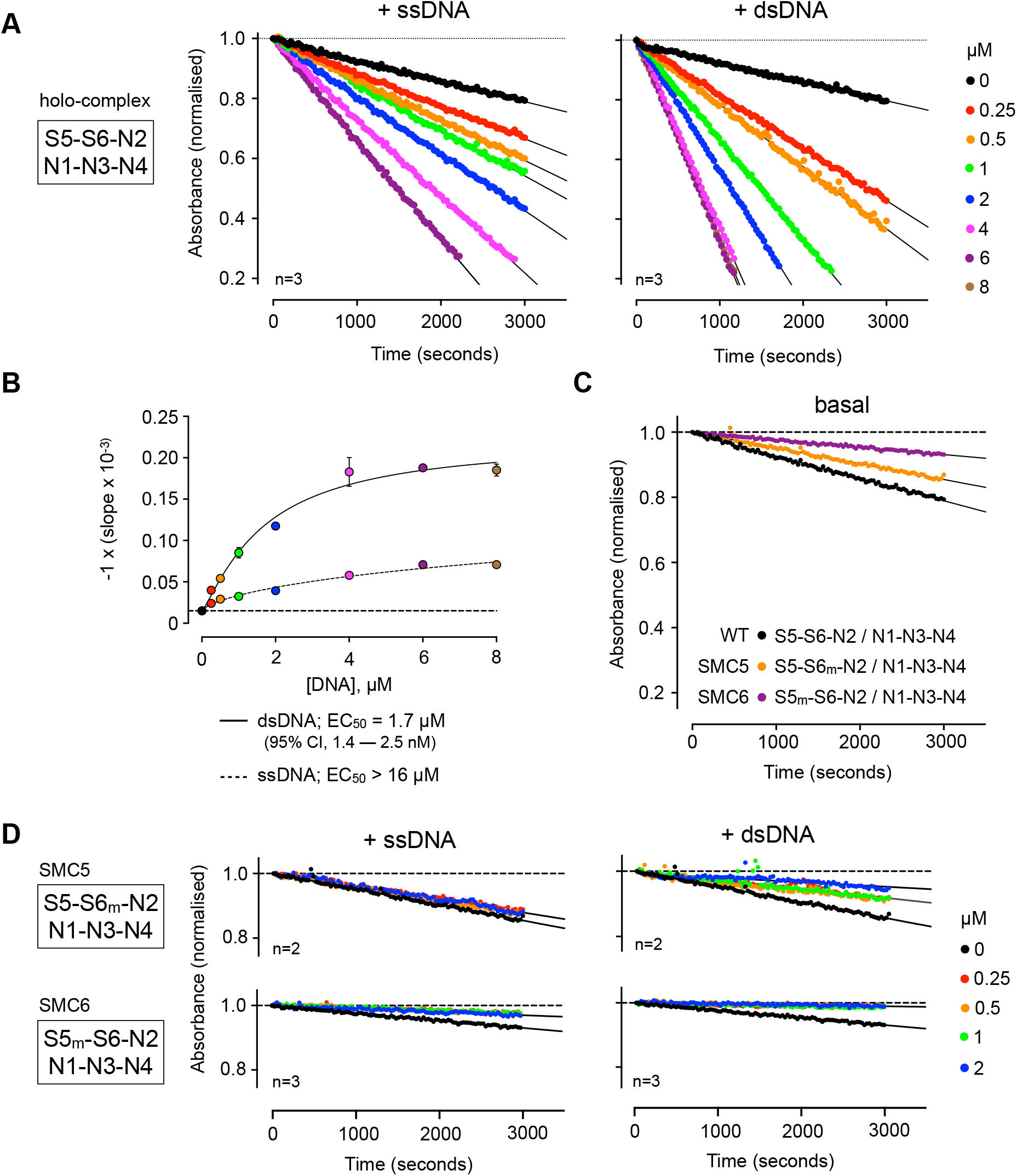
ATP turnover is preferentially stimulated by addition of double-stranded DNA. (**A**) Titration of single-stranded or double-stranded DNA into the Smc5/6 holo-complex, stimulates ATP turnover in a NADH-coupled regenerative ATPase assay. Least-squares fitting of a straight-line equation to experimental data provides a ‘slope’ parameter for each titration point (change in absorbance over time; solid black line). (**B**) Estimation of EC50 for titrations, by fitting of a stimulatory dose-response model to experimental data by non-linear regression. 95% CI = confidence interval. (**C**) The basal level of ATP turnover by SMC5 is greater than SMC6, as judged by experiments using purified holo-complexes containing a single Walker B mutation in one or another SMC subunit. (**D**) The stimulation of ATPase activity generated by addition of nucleic acid is lost in reconstituted holo-complexes that contain a single Walker B mutation in one or another SMC subunit. 5_m_ = SMC5-E1015Q, 6_m_ = SMC6-E1048Q.

Recently published data has demonstrated that the ability of the individual SMC proteins to turn over ATP, within either condensin or cohesin, is not equivalent^37,38^. To confirm if the same held true for Smc5/6, we purified complexes where just Smc5 or Smc6 harboured the aforementioned E-to-Q mutations and measured activity. With introduction of either mutation, a reduced basal level of activity was evident, but this was approximately 2-fold higher when Smc6 was disabled as compared to the Smc5 mutant (**Figure 4B**). We then repeated the titrations of ssDNA / dsDNA into each of the mutated complexes, however this time observing no stimulation of ATPase activity in either case. Instead, addition of nucleic acid resulted in a net decrease from basal levels of activity, with dsDNA having the greater effect (**Figure 4C**).

### An inhibitory effect of adding Nse5/6

We next added increasing concentrations of purified recombinant Nse5/6 to the holo-complex (reaching 1.5 molar equivalents) and determined the effect on ATPase activity in both the absence and presence of dsDNA (**Figure 5**). Addition of Nse5/6 to the reconstituted holo-complex inhibited its ability to turn over ATP, even in the presence of two different stimulatory concentrations of dsDNA (2 and 10 μM). Plotting slope (decrease in absorbance per second) versus molar equivalents of Nse5/6 revealed that generation of a 1:1 stoichiometric complex was sufficient to fully inhibit ATP-turnover (**Figure 5**).

**Figure 5.**
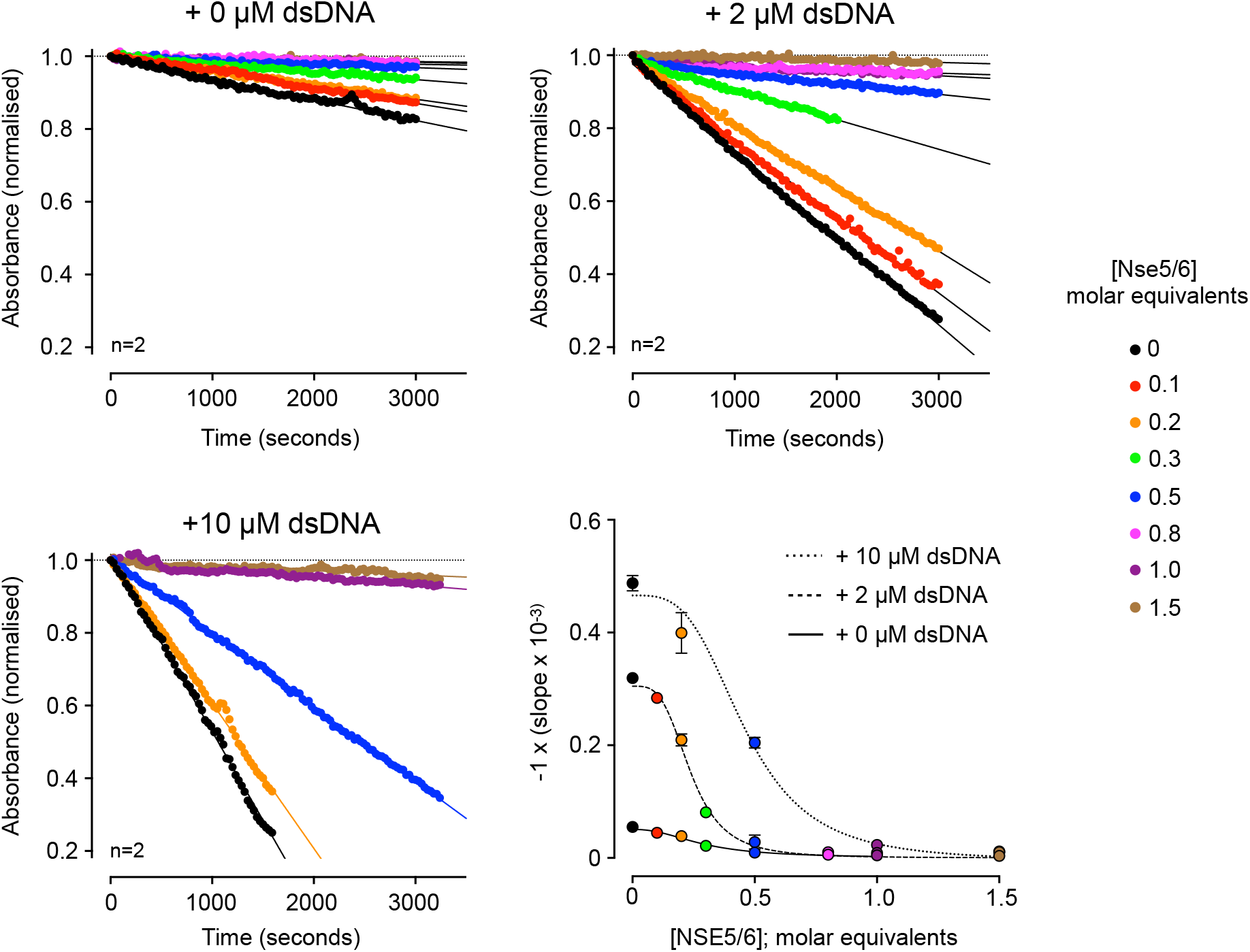
Addition of Nse5/6 to the Smc5/6 holo-complex inhibits ATPase activity. Titration of Nse5/6 inhibits the ability of the Smc5/6 holo-complex to turn over ATP, even in the presence of stimulating concentrations of double-stranded DNA (2 and 10 μM); as shown by experiments carried out in an NADH-coupled regenerative ATPase assay. A plot showing change in absorbance with time (slope) against molar equivalents of Nse5/6 added, reveals that formation of a 1:1 equimolar complex between Nse5/6 and Smc5/6 holo-complex is sufficient to prevent ATP-hydrolysis. Fitted curves are for visualisation purposes only.

### DNA-binding activity of reconstituted complexes

To help define the mechanism of ATPase inhibition, we investigated the possibility that Nse5/6 might directly disrupt or compete with the ability of the holo-complex to bind to dsDNA. We first performed a series of control experiments, confirming the ability of the purified holo-complex to bind a fluorescently labelled 48nt dsDNA hairpin using electrophoretic mobility shift assays (EMSA) (**Figure 6A**). On titration of the holo-complex, an initial complex was formed (I) which was then super-shifted into a second, slightly slower migrating species (II) at higher protein concentrations. Quantification of free and bound states allowed an overall dissociation constant *K*_d_ of ~220 nM to be estimated for the interaction (**Figure 6B**). At this time, we also took the opportunity to test the DNA-binding capability of the S5-S6-N2 ‘core-complex’ under the same set of experimental conditions. Whilst some low-level aggregation / precipitation of the complex was evident at higher concentrations (manifested as fluorescent material stuck in the loading pockets of the agarose gel) there was no compelling evidence for an interaction with the hairpin (**Figure 6C**). Similarly, Nse5/6 also did not bind the hairpin over a more extensive concentration range (**Figure 6D**).

**Figure 6.**
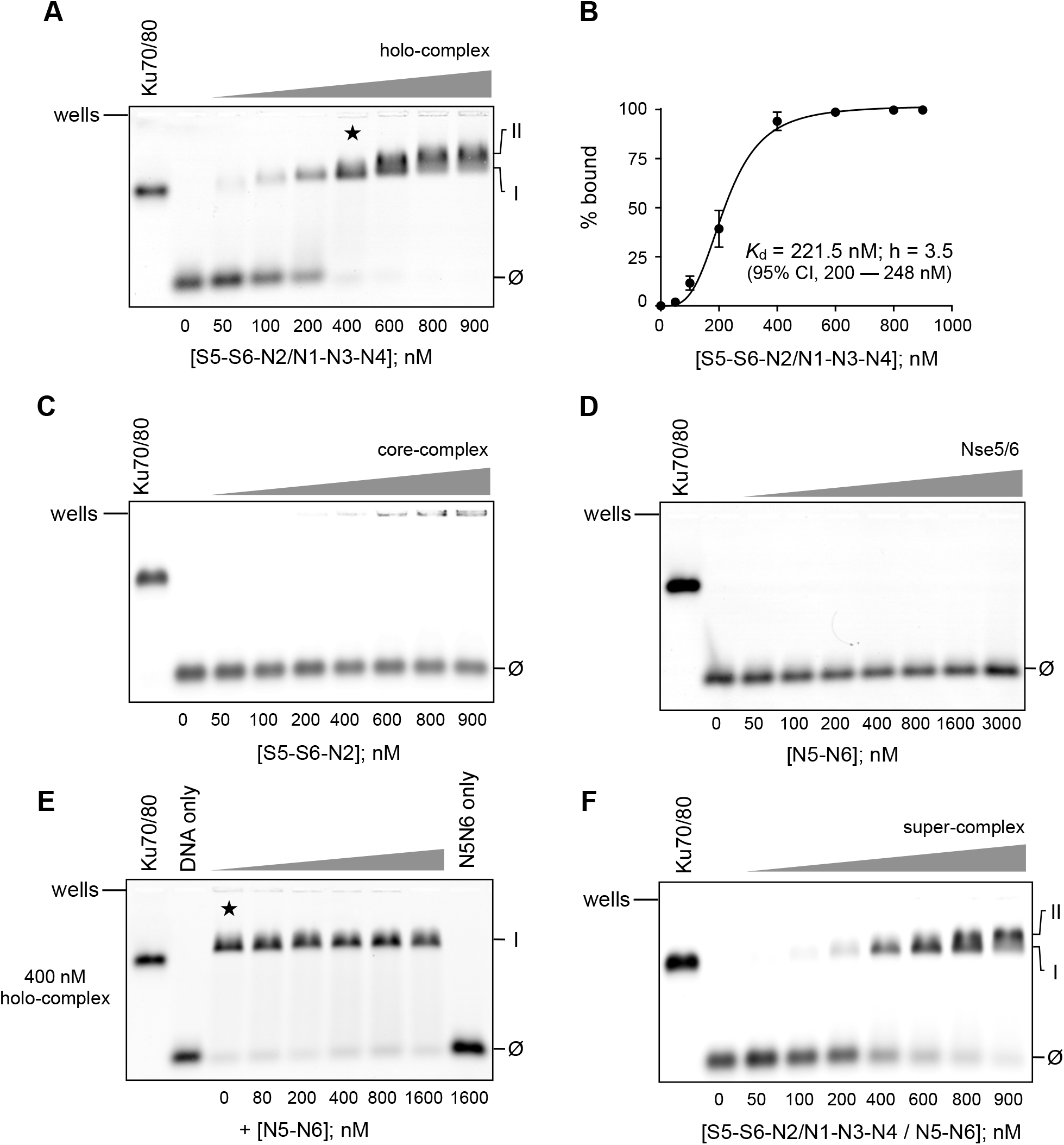
Nse5/6 does not perturb the ability of the Smc5/6 holo-complex to bind dsDNA. (**A**) Electrophoretic mobility shift assay (EMSA) showing binding of the Smc5/6 holo-complex to a fluorescently labelled double-stranded DNA hairpin. An initial complex is formed (labelled as ‘I’) which at higher concentrations is then super-shifted into a second, slightly slower migrating complex (‘II’). (**B**) Quantification of free and bound species allows a dissociation constant (*K*d) of ~220 nM to be estimated, with a hill-slope parameter (h) of 3.5. Errors bars represent 1 standard deviation across 3 experimental repeats. 95% CI = confidence interval for value of *K*d obtained by least-squares fitting of a binding model to the experimental data. (**C**) The ‘core’ complex comprised of just Smc5, Smc6 and Nse2 is not able to bind to the dsDNA harpin, as judged by EMSA. At higher concentrations, some aggregated or precipitated fluorescent material can, however, be visualised in the wells of the agarose gel. (**D**) Purified Nse5/6 heterodimer does not bind to the dsDNA hairpin, as judged by EMSA. (**E**) In a competition experiment, increasing concentrations of Nse5/6 were added to a pre-formed complex made between the Smc5/6 holo-complex and the dsDNA hairpin (400 nM, as marked by the five-pointed star in panel A). However, no disruption to the ability of the holo-complex to bind dsDNA was evident. (**F**) The Nse5/6 containing ‘super-complex’ is still capable of binding to the dsDNA hairpin, as judged by EMSA. Ku70/80 heterodimer is included as a positive control in each experiment. Ø marks the migration position of the unbound dsDNA hairpin.

A competition experiment (**Figure 6E**) in which increasing amounts of Nse5/6 were added to a pre-formed complex — generated under conditions where the majority of the DNA hairpin is bound (400 nM) — showed no disruption of dsDNA binding by the Smc5/6 holo-complex, even in the presence of excess Nse5/6. We then repeated the EMSA using the purified ‘super-complex’ containing Nse5/6 (see next section), seeing no significant perturbation or major changes to the ability of the complex to bind the dsDNA hairpin (**Figure 6F**).

### Visualisation of the NSE5/6-bound SMC5/6 complex

Initial negative stain experiments revealed a mix of unliganded (holo-complex) and liganded (Nse5/6-containing) states in the applied sample (data not shown). Inclusion of an additional purification step (C.tag affinity resin) allowed selective enrichment of the Nse5/6-containing ‘super-complex’ and greatly improved sample homogeneity (**Materials and Methods, Figure 7A).** After stabilisation by crosslinking and application to a size exclusion chromatography column (**Materials and Methods**) we verified that Nse5/6 was present in the fraction selected for negative-stain experiments by western blot, using an HRP-conjugated nanobody that recognises the C.tag epitope on Nse5 (**Materials and Methods**, **Figure 7B**, lane X2). We then used the same visualisation / focussed refinement strategy as before to produce a 3D model (**Figure 7, panels C to E, Table 1**).

**Figure 7.**
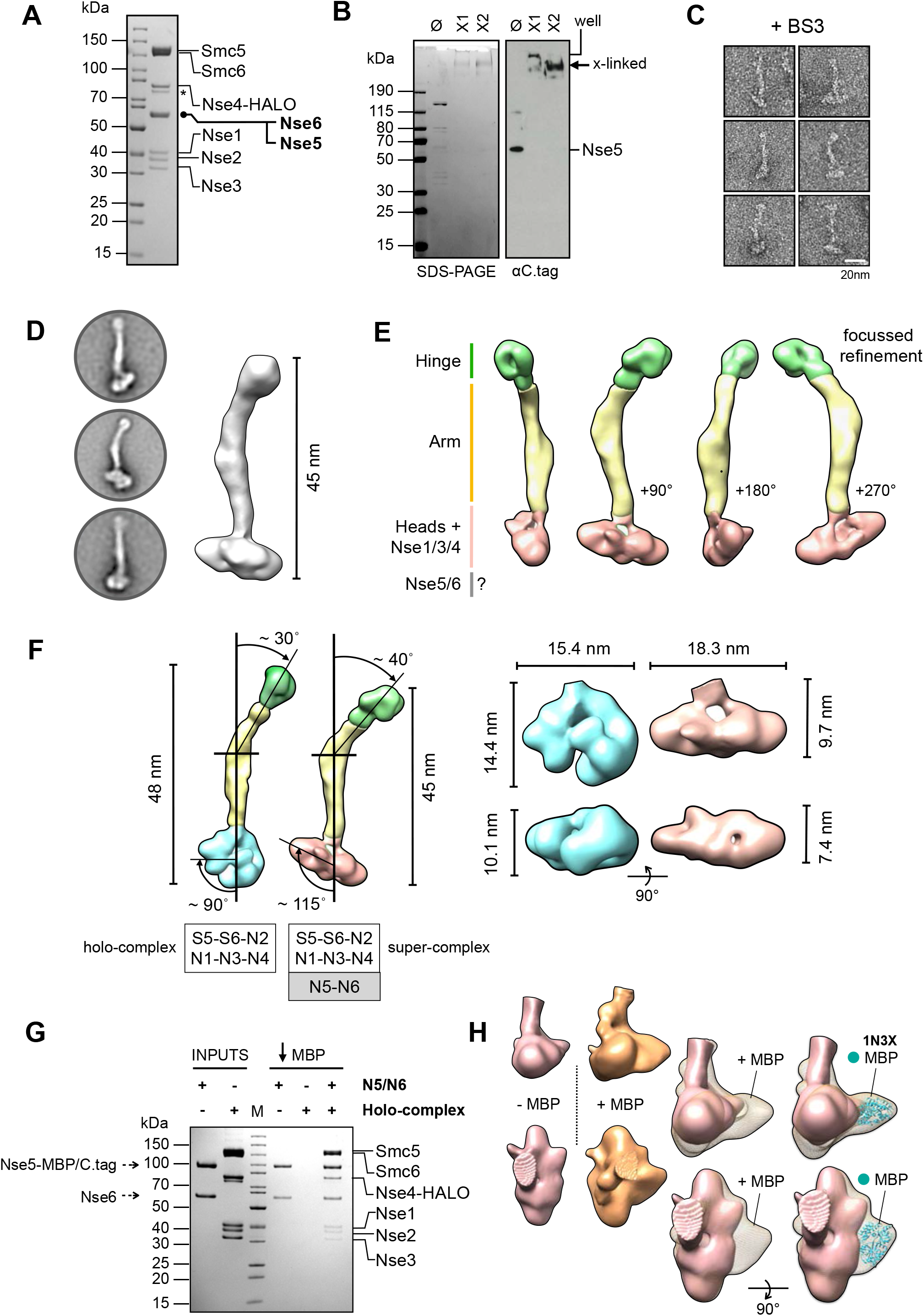
Addition of the NSE5/6 heterodimer to the Smc5/6 holo-complex. (**A**) Colloidal-blue stained SDS-PAGE gel of the purified Nse5/6-containing ‘super-complex’. (**B**) SDS-PAGE gel and associated western blot showing the super-complex before and after crosslinking with BS3. Nse5, which carries a C-terminal C.tag epitope is detected by a nanobody coupled to horse-radish peroxidase (αC.tag). Ø = uncrosslinked. X1 and X2 = successive elution fractions from a size exclusion chromatography column. (**C**) Representative images of particles, taken from micrographs of uranyl acetate negative-stained super-complex. (**D**) Representative 2D class averages and initial 3D reconstruction. (**E**) Segmented 3D reconstruction resulting from focussed refinement strategy (**F**) Side-by-side comparison of 3D volumes for both the Smc5/6 holo-complex and Nse5/6-containing ‘super-complex’. Using a section of the ‘arm’ closest to the ‘head-end’ of the complex to define a vertical axis, allows changes in overall conformation to be described by the indicated angles (**G**) Colloidal-blue stained SDS-PAGE for a pull-down experiment on amylose chromatography beads, demonstrating that C-terminal fusion of Nse5 to Maltose-binding protein (MBP) does not affect its ability to bind to the holo-complex, in the context of the Nse5/6 heterodimer. (**H**) Side-by-side comparison of 3D volumes obtained by focussed refinement, for the head-end of super-complexes containing either Nse5-C.tag (coloured pink) or Nse5-MBP/C.tag (orange). The additional protruding lobe of density is consistent the dimensions of the X-ray crystal structure of *E. coli* MBP (PDB ID: 1N3X).

A change in the conformation of the complex was immediately evident (**Figure 7E and 7F**). By using the lower section of the ‘arms’ as a vertical reference axis, we could measure an apparent increase in the tilt angle of the hinge, from ~30° in the holo-complex to ~40° in the Nse5/6-containing super-complex, which also manifests as an apparent reduction in its overall length to around 45 nm. A more drastic change was observed at the head end of the complex, with a broadening of its overall width combined with a compaction in height (**Figure 7F, right**). Another angle, used to relate the head-end of the complex to the reference axis, also changes from ~90° in the holo-complex to ~115° in the super-complex (**Figure 6F**).

It was not possible, however, to unambiguously identity a region of density that matched the globular shape of the isolated Nse5/6 heterodimer, that we had also visualised by negative stain (**Supplementary Figure 3A**). We therefore remade our Nse5/6 baculovirus, engineering a C-terminal fusion between Nse5 and Maltose Binding Protein (that also carried the C.tag epitope; MBP / C.tag). We confirmed that the protein-fusion was still able to bind to the holo-complex by a pull-down experiment on amylose chromatography resin (**Figure 7G**), and then once again selectively purified, stabilised and visualised the resulting complex (**Table 1**).

No major differences or changes to either the ‘arm’ or the ‘hinge’ regions were evident but focussed 3D refinement revealed an increased volume at the ‘head’ end of the complex (**Supplementary Figure 3B**). Side-by-side comparison of the two Nse5/6-containing complexes allowed identification of a lobe of additional density, compatible with the expected volume for MBP, thus serving to localise bound Nse5/6 to the ‘head’ end of the Smc5/6 complex (**Figure 7H**).

## DISCUSSION

We have reconstituted a set of defined budding yeast Smc5/6 complexes using recombinant proteins expressed in insect cells. By taking this ‘bottom up’ approach we are able to carefully control and examine different subunit compositions and stoichiometries, plus the ability to introduce mutations that would otherwise be incompatible with viability of the native host.

Biochemical experiments with our reconstituted complexes reveal that the 6-component holo-complex (formed of Smc5, Smc6, Nse1, Nse2, Nse3 and Nse4) is capable of hydrolysing ATP and binding to a short dsDNA hairpin, whereas the minimal 3-component ‘core’ complex is not (Smc5, Smc6, Nse2). We observe that ATP hydrolysis by the holo-complex is strongly stimulated by the addition of a dsDNA substate to reaction mixes, and that the two ATPase activities of Smc5/6 are not equivalent — consistent with other studies of heterodimeric SMC-complexes^37–42^— the basal hydrolysis rate of Smc5 being approximately 2-fold higher than that of Smc6. DNA-dependent stimulation of ATPase activity is, however, wholly dependent on functional active sites in both SMC proteins, as it is precluded when Walker B, E-to-Q mutations are introduced into one or other subunit. Taken as a whole, our data confirm that there is a high degree of functional coupling between the two active sites of Smc5/6 that is directly linked to dsDNA binding, and which is dependent on the Nse1-Nse3-Nse4 subcomplex being bound to the ‘head’ end of the Smc5/6 complex.

Visualisation of our reconstituted Smc5/6 complexes by negative stain electron microscopy reveal that they adopt a predominantly rod-like or ‘extended’ conformation, with no acute bend at an ‘elbow’ as has been recently observed for *E. coli* MukBEF as well as the cohesin and condensin complexes^43–49^. However, the appearance of individual particles, plus the requirement for focussed refinement of 3D models, indicates that there is still a degree of conformational flexibility (**Figures 1C, 1D and 7C**). We estimate the overall length of the holo-complex to be 49 nm, with an arm length of approximately 27 nm; distances compatible with a computational study that has indicated an overall shortening of the ‘arm’ length within the Smc5/6-family, when compared to cohesin and condensin complexes^50^.

We have located the position of Nse5/6 binding to the ‘head-end’ of the complex (**Figure 7H**), in contrast to a previous study that indicated binding to the hinge^32^. Pleasingly, this now serves to unify the architecture of the *S. cerevisiae* Smc5/6 complex with its *S. pombe* orthologue^33^. Concomitant with association of Nse5/6 is a significant change in conformation at the head-end of the complex, plus an overall reduction in its length to ~45 nm (**Figure 7**).

Nse5/6 has been identified as a factor required for recruitment of Smc5/6 to collapsed replication forks and for recruitment and/or retention of the complex at defined chromatin sites, which include highly repetitive sequences such as centrosomes, telomeres and the ribosomal DNA array^51,52^. It has been proposed to function in a manner similar to the Scc2-Scc4 cohesin loader complex^22^, but it is worth noting here that Scc2-Scc4 has been shown to stimulate (rather than inhibit) the ATPase activity of cohesin and is capable of binding directly to DNA^53,54^. Nse5/6 therefore cannot work in exactly the same manner, as we show here that it serves to inhibit ATPase activity and has no intrinsic ability to bind DNA. So, whilst Nse5/6 can act as an intermediary, bringing Smc5/6 to sites of replication stress through its own interaction with Rtt107 (Brc1 in *S. pombe)*, which itself binds to γH2A via its C-terminal BRCT-pair^55–57^, it is still not clear how it works to promote chromatin-binding and retention.

Another requirement for stable chromatin association of the Smc5/6 complex is the ability of the Nse3 subunit to bind dsDNA, as revealed by ChIP experiments carried out in fission yeast^58^. Whilst mutations that fully disrupt DNA-binding are lethal when introduced into *S. pombe*, those that act to reduce DNA-binding are tolerated. Strains carrying such hypomorphic mutations have reduced viability, display sensitivity to a range of different genotoxic agents, and have a global reduction in the levels of Smc5/6 precipitated at a range of different chromatin loci^58^. Recent single-molecule live cell imaging experiments, also carried out in fission yeast, show that mutations which perturb the ability of the Smc5/6 complex to turn over ATP result in a decreased level of chromatin association, as does introduction of the hypomorphic nse3-R254E allele identified by Zabrady *et al.* (ref. 58), that disrupts (but does not abolish) dsDNA binding. Furthermore, deletion of the gene encoding Nse6 results in an almost complete loss of chromatin associated Smc5/6^25^.

A complex, likely allosteric, mechanism that involves the interplay of dsDNA-binding and ATP-hydrolysis by the Smc5/6 holo-complex, plus binding (and release?) of the Nse5/6 heterodimer appears to be at play, serving to promote chromatin-binding and/or retention of the Smc5/6 complex. Our discovery that Nse5/6 negatively regulates ATP-hydrolysis by the *S. cerevisiae* Smc5/6 complex provides a new and potentially important piece of information.

A caveat of our study is that we do not know the nucleotide-bound status of our purified complexes and therefore cannot unambiguously assign 3D models to defined states i.e., apo, ATP- or ADP-bound. However, when present at 1:1 stoichiometry (relative to the holo-complex) binding of Nse5/6 prevents all ATP-hydrolysis, at both basal and DNA-stimulated levels. Invoking Occam’s razor, the simplest explanation is that binding of Nse5/6 blocks the ability of the two head domains of Smc5/6 to engage productively with each other: a hypothesis compatible at least with its observed position of binding. However, to fully understand the molecular details of the interface between Nse5/6 and the Smc5/6 complex, and the resultant set of conformational changes that appear to underpin function, obtaining structural data at higher resolution is now a desirable goal.

## General

The authors would like to thank Prof. Laurence Pearl (Uni. of Sussex/ICR) and Prof. Antony Carr (Uni. of Sussex) for constructive criticism and proof-reading of this manuscript. They would also like to thank Dr. Neil Kad (Uni. of Kent) for providing details of the NADH-coupled regenerative ATPase assay.

## Author Contributions

Conceptualisation: JMM, AWO; Methodology: STH, PS, FB, EM, JMM, AWO; Investigation: STT, PS, LZ, FB, AWO; Writing — Original Draft: STH, AWO; Writing — Review and Editing: STH, PS, FB, EM, JMM, AWO; Visualisation: AWO; Supervision: PS, FB, EM, JMM, AWO; Funding Acquisition: JMM, AWO.

## MATERIALS AND METHODS

### Expression constructs

#### Smc5/6

Synthetic genes codon-optimised for expression in *Spodoptera frugiperda*, for each of the proteins forming the *Saccharomyces cerevisiae* Smc5/6 complex, were purchased from GeneArt [ThermoFisher Scientific, Loughborough, UK]. In each case, the coding sequence was subcloned into the vector pLIB of the biGBac system at the BamHI and HindIII sites within the multiple cloning site^59^. With the exception of Nse3, amino acids encoding in-frame affinity/epitope tags were added at either the start or end of the coding sequence.

Expression constructs were then generated via PCR amplification / Gibson Assembly reactions following the procedures and protocols published in ref. 59.

1. [S5S6N2]: pBIG1a, containing Smc5-FLAG, Smc6-AVI-HIS and HA-Nse2
2. [N1N3N4]: pBIG1b, containing HIS-TEV-Nse1, Nse3, Nse4-HALO-Myc
3. [N5N6]: pBIG1c, Nse5-C.tag, HIS-3C-Nse6
4. [S5S6N2-N1N3N4]; pBIG2ab, generated by combining constructs 1 and 2
5. [N5MBPN6]: pBIG1c, Nse5-MBP-C.tag, HIS-3C-Nse6

AVI-HIS: combined Avi^60^ + His_6_affinity tag with spacer; GLNDIFEAQKIEWHEGSASGHHHHHH

C.tag: GAAEPEA^61^
FLAG: DYKDDDDK
HALO: HaloTag; modified haloalkane dehalogenase^62^
HA: YPYDVPDYA
HIS: HHHHHH
MBP: Maltose-binding protein MYC: EQKLISEEEDL
TEV: Tobacco Etch Protease cleavage site; ENLYFQG
3C: Human Rhinovirus 3C protease cleavage site; LEVLFQGP

#### C.tag nanobody

A synthetic gene, codon-optimised for expression in *E.coli*, was purchased from GeneArt. The coding sequence was subcloned into the NcoI and XhoI restrictions sites of the expression vector pCDF-1b [Merck KGaA, Darmstadt, Germany] in-frame with a C-terminal non-cleavable 6xHis affinity tag.

### Expression in insect cells

Recombinant viruses were generated using the Bac-to-Bac Baculovirus Expression System [ThermoFisher Scientific]. All expression was carried out in *Sf*9 insect cells, as suspension culture in 2 L bottles containing 500 ml Insect-Xpress media [Lonza Bioscience, Slough, UK] supplemented with penicillin and streptomycin. Cells at a density of 2 x 10^6^ cells/ml were infected with the appropriate viral stock at a multiplicity-of-infection of 2, then grown in an orbital shaking incubator set at 27 °C and 150 rpm, for a period of 72 hours. Cells were harvested by centrifugation at 1500 x *g* for 10 minutes and the resulting cell pellet stored at −80°C until required.

### Expression in *E. coli*

*E. coli* strain BL21(DE3) [New England Biolabs, Hitchin, UK)] was transformed with the C.tag-nanobody expression vector, selected by plating on LB-agar plates supplemented with 50 μg/ml spectinomycin. Transformed cells were used to inoculate a 250 ml conical flask containing 50 ml Turbo-broth [Molecular Dimensions, Sheffield, UK] supplemented with antibiotic as before. The culture was grown at 37°C in an orbital-shaker incubator until an OD_600_ of 1.5 was reached, it was then stored at 4°C overnight. 12 ml from this ‘starter culture’ was then used to inoculate a 2L conical flask containing 1L of Turbo-broth media, supplemented with antibiotic as before. This culture was incubated at 37°C in an orbital-shaker incubator, as before, until an OD_600_ of 1.5 was reached. The flask containing the culture was chilled on a bed of ice for a period of 1 hour before induction of recombinant protein expression by the addition of 0.2 mM isopropyl β-D-1-thiogalactopyranoside [Generon Ltd., Slough, UK]. Cultures were then incubated at a lower temperature of 20°C in an orbital-shaker incubator for a period of ~16 hours, before harvesting of cells by centrifugation at 4000 x *g* for 20 minutes. The resultant cell pellet was stored at −80°C until required.

### Protein purification

#### Buffer Composition

A: 50 mM HEPES.NaOH pH 7.5, 250 mM NaCl, 10 mM imidazole, 0.5 mM TCEP
B: 50 mM HEPES.NaOH pH 7.5, 250 mM NaCl, 300 mM imidazole, 0.5 mM TCEP
C: 20 mM HEPES.NaOH pH 7.5, 100 mM NaCl, 0.5 mM TCEP
D: 20 mM HEPES.NaOH pH 7.5, 150 mM NaCl, 0.5 mM TCEP
E: 20 mM HEPES.NaOH pH 7.5, 750 mM NaCl, 0.5 mM TCEP
F: 20 mM HEPES.NaOH pH 7.5, 0.5 mM TCEP
G: 20 mM MES.NaOH pH 6.5, 100 mM NaCl, 0.5 mM TCEP
H: 20 mM MES.NaOH pH 6.5, 1000 mM NaCl, 0.5 mM TCEP
I: 20 mM HEPES.NaOH pH 7.5, 250 mM NaCl, 0.5 mM TCEP

#### Peptides

3 x FLAG: MDYKDHDGDYKDHDIDYKDDDDK

SEPEA: SEPEA

#### Smc5/6 complexes

The cell pellet from 1L of Sf9 cell culture was resuspended on ice, in BUFFER A supplemented with protease inhibitors [Roche, Burgess Hill, UK]. Cells were lysed though a combination of the thawing process and hand-homogenisation, and insoluble material removed by high-speed centrifugation at 40,000 *x g* for a period of 1 hour at 4°C. A protamine sulphate precipitation step was also included to remove excess nucleic acid. The soluble supernatant was then filtered through a 5 μm filter [Sartorius Stedim, Epsom, UK] then applied to a batch/gravity column containing 2 ml Amintra CoHIS resin [Expedeon, Over, UK], pre-equilibrated in BUFFER A. The resin was washed with 10 column volumes (CV) of BUFFER A, then the retained protein eluted by application of 5 CV of BUFFER B. The eluate from this step was diluted 2-fold with BUFFER C in order to reduce the overall NaCl and imidazole concentration, and then incubated with 0.5 ml ANTI-FLAG M2 Affinity Gel [Sigma-Aldrich Company Ltd, Gillingham, UK] pre-equilibrated in BUFFER D for a period of 1 hour with rolling at 4°C. The resin was collected in a gravity flow column and then washed with 5 CV BUFFER D, with bound protein eluted by application of the same buffer containing 0.2 mg/ml 3xFLAG peptide [Peptide Protein Research Ltd, Waltham, UK].

Pooled fractions were loaded onto a 1 ml Heparin column [Cytiva, UK] to concentrate the complex and to remove any peptide from the previous chromatography step. The column was washed with 5 CV BUFFER D, before elution of retained protein with BUFFER E.

Finally, the eluate was applied to a Superose 6 Increase GL size exclusion chromatography (GE Healthcare Life Sciences, Little Chalfont, UK) pre-equilibrated in BUFFER D. Throughout the purification procedure, samples were analysed by SDS-PAGE in order to monitor yield and purity.

Enrichment of complexes containing the C.tag epitope was achieved by inclusion of the following chromatography step, placed after the initial capture by IMAC and dilution with BUFFER C. The diluted eluate was incubated with 0.3 ml CaptureSelect C-tag XL Affinity Matrix [ThermoFisher Scientific] pre-equilibrated in BUFFER D for a period of 1 hour with rolling at 4°C. The resin was collected in a gravity flow column and then washed with 5 CV BUFFER D, with bound protein eluted by application of the same buffer containing 2 mM ‘SEPEA’ peptide [Peptide Protein Research]. Samples were then applied sequentially to the heparin and size exclusion chromatography columns as before.

#### C-tag nanobody

The cell pellet from 1 L of cell culture was resuspended in BUFFER A supplemented with protease inhibitors [Roche]. Cells were disrupted, on ice, by sonication at 40 % amplitude in bursts of 5 seconds on and 20 seconds off, for a total of 5 minutes [Vibra-Cell VCX500 Ultrasonic Processor], with insoluble material removed by centrifugation. The resulting supernatant was filtered through a 5 μm sterile syringe filter [Sartorius Stedim] and then loaded onto a 5 ml HiTrap TALON crude column [Cytiva, Little Chalfont, UK] pre-equilibrated in BUFFER A. Unbound material was removed by washing with 5 CV BUFFER A, before retained protein was eluted by application of BUFFER B.

Pooled fractions were concentrated using Vivaspin 20 (10 000 MWCO) centrifugal concentrators [Sartorius] before buffer-exchange into BUFFER F, and then application to a 5 ml HiTrap SP FF cation exchange column [Cytiva] pre-equilibrated in the same buffer. Unbound material was removed by washing with 5 CV BUFFER F, before eluting the retained protein with a linear salt gradient made with BUFFER H over ~12 CV. Fractions containing the desired protein were pooled and concentrated to a final volume of 5 ml using centrifugal concentrators as before, then applied to Superdex 75 26/600 size exclusion column [Cytiva] pre-equilibrated with BUFFER I as the final purification step. Fractions containing the protein complex were identified by SDS-PAGE, pooled and concentrated as before to a final concentration of 6.5 mg/ml. The purified protein was then flash frozen in liquid N_2_ and stored at −80°C until required.

### HRP Conjugation

The C.tag-nanobody was conjugated to horseradish peroxidase, using HRP Conjugation Kit – Lightning-Link from Abcam [Ab102890, Cambridge, UK] as per manufacturer’s instructions. The resulting C-tag-nanobody-HRP conjugate was stored at 4°C until required.

### Cross-linking

Complexes used in negative stain experiments were cross-linked either with 0.1% glutaldahyde [Agar Scientific Ltd, Stansted, UK] for a period of 10 minutes at room temperature or overnight (~15 hours) at 4°C with 1 mM BS3 (bis(sulfosuccinimidyl)suberate) [Fisher Scientific UK Ltd., Loughborough, UK]. Reactions were stopped by the addition of 50 mM Tris pH 7.5. Samples were then concentrated using Vivaspin 6 (50 000 MWCO) centrifugal concentrators [Sartorius Stedim] to a volume of 0.5 ml before being applied to a Superose 6 Increase GL size exclusion chromatography [Cytiva] pre-equilibrated in BUFFER D, as the final purification step. Samples were used immediately for both assays and negative stain grid preparation.

### Antibodies (Western blot)

#### Primary antibodies

Anti-His_6_; mouse monoclonal at 1:10,000 dilution (631212, Merck)
Anti-FLAG M2; mouse monoclonal at 1:10,000 dilution (F3165, Merck)
HA-Tag antibody; mouse monoclonal at 1:10,000 dilution (sc-7392, Santa Cruz Biotechnology)
Anti-Myc tag antibody; mouse monoclonal at 1:10,000 dilution (ab32, Abcam)
CaptureSelect Biotin Anti-C-tag conjugate; camelid antibody fragment at 1:10,000 dilution (7103252100, ThermoFisher)
C.tag HRP-conjugate; nanobody at 1:20,000 (in house)

#### Secondary antibodies

Amersham ECL mouse IgG HRP-linked whole Ab; sheep polyclonal at 1:10,000 dilution (NA931V, Cytiva)

Streptavidin-Horseradish Peroxidase Conjugate at 1:10,000 dilution (RPN1231, Cytiva)

### Biochemical and Biophysical Assays

#### Oligonucleotide Sequences

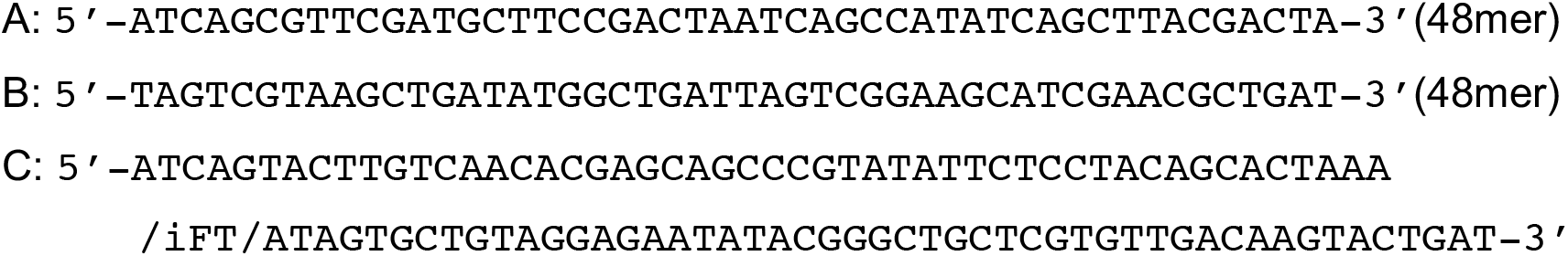

Where iFT = fluorescein attached to position 5 of the thymine ring by a 6-carbon spacer. All experiments with ssDNA use oligonucleotide A. Those with dsDNA use a DNA duplex formed by annealing oligonucleotide A with B, with the exception of DNA-binding experiments (EMSA) that used self-annealed oligonucleotide C.

Purified unlabelled DNA oligonucleotides used in NADH-coupled ATPase assays were purchased from Merck. Purified fluorescein-labelled DNA oligonucleotides used in EMSA experiments were purchased from Integrated DNA Technologies [Leuven, Belgium].

### NADH-coupled regenerative ATPase assay

Methodology is based on that previously reported by Barnett and Kad ^36^. All reagents were purchased from Merck. Assays were performed in 96 well UV-transparent plates at 25°C in BUFFER D. 143 nM of each complex was incubated with a 1/60 dilution of Pyruvate Kinase / Lactic dehydrogenase (PK/LDH; P0294), 0.42 mM Phospoenolpyryvate (PEP;10108294001), 0.83 mM ATP-MgCl_2_ and 0.176 mM beta-Nicotinamide adenine dinucleotide hydrogen (NADH;10128023001) in a total volume of 120 μl. Absorbance at 340 nm was then measured every 30 seconds for a total of 5400 seconds using a CLARIOstar multimode plate reader [BMG Labtech GmbH, Baden-Württemberg, Germany]. Data were processed and analysed using GraphPad Prism [v. 9.0, GraphPad Software LLC, San Diego, US].

### EMSA

Self-annealed oligonucleotide C (at a concentration of 200 nM) was pre-incubated with the indicated dilution series of each complex, in BUFFER D for a period of 30 minutes at 4°C. NativePAGE 4x loading buffer [Fisher Scientific, Loughborough, UK] was then added to each sample, before being applied to 0.8% w/v tris-borate-EDTA (TBE)-agarose gel. Electrophoresis was carried out by application of 50V for a period of 3 hours at 4°C, in 0.5 % v/v TBE (Novex TBE Running Buffer, Fisher Scientific). Separated species were visualised using a Fuji FLA-5100 Fluorescent Image Analyser, using excitation with a 473 nm laser.

### MBP Pull-down

Pull-down experiments were performed in BUFFER D. 300 μl of the indicated complex at 1 μM was incubated with 100 μl Amylose Resin [New England Biolabs, Hitchin, UK] for a period of 1 hour at 4 °C with rolling/agitation. The resin was then collected by centrifugation at 100 x *g* for 1 minute at 4 °C, in a 500 μl Corning Costar Spin-X Plastic Centrifuge Tube [0.45 μm, Merck, Gillingham, UK]. The resin was washed 3 times by application of 500 μl BUFFER D, before bound material was eluted through application of BUFFER D supplemented with 20mM maltose.

### Uranyl Acetate Negative Stain Transmission Electron Microscopy

#### Grid preparation

Copper grids (continuous carbon film on 300 mesh, Agar Scientific Ltd, Stansted, UK) were treated by glow discharge for 60 to 90 secs at 15 mA (PELCO EasyGlow). 3 μl of sample was placed onto the glow-discharged grids for a period of 1 min before excess liquid was removed by gently blotting with filter paper. Grids were then washed 3 times with either ultrapure water or BUFFER D, then stained with 2% uranyl acetate for 30 secs and dried thoroughly with filter paper.

### Data collection

S5-S6-N2/N1-N3-N4 dataset was collected on a Tecnai TF20 transmission electron microscope, equipped with a TemCam F416 CMOS camera at the Institute of Cancer Research (London, UK) at an excitation voltage of 200 kV. In total, 416 images were collected at 50,000x magnification and a pixel size of 1.732 Å.

Nse5/Nse6-containing datasets were collected in house, on a JEOL JEM-1400-plus transmission electron microscope, at 120 kV, equipped with a OneView camera (Gatan, Inc). Acquisition was performed at 25 fps with 1 sec integration time and drift correction performed automatically using the Gatan Microscopy suite (GMS3, Gatan, Inc). In total, 234 and 102 images were collected at 60,000x magnification and a pixel size of 1.8 Å.

### Data processing

Micrographs were processed with RELION (v3.1; ^63,64^. Individual particles were picked manually and extracted using box sizes of 352 and 328 pixels for S5/S6/N2-N1/N3/N4 and Nse5/6-containing complexes respectively. 2D classification was used an initial ‘polishing’ step, to select particles to take forward into 3D classification. Particles that converged into a single class, were then used in 3D refinement. A mask diameter of 550 Å was used in both 2D and 3D classification steps. Masks for the ‘Head’, ‘Arm’ and ‘Hinge’ regions of each complex were generated from 3D volumes using Chimera (v1.14; ^65^, and then used for 3D refinement to improve particle alignment within these regions. See Table I for additional information.

## Figures

Molecular images were generated using either PyMOL ^66^ or Chimera ^65^.

**Figure S1.**
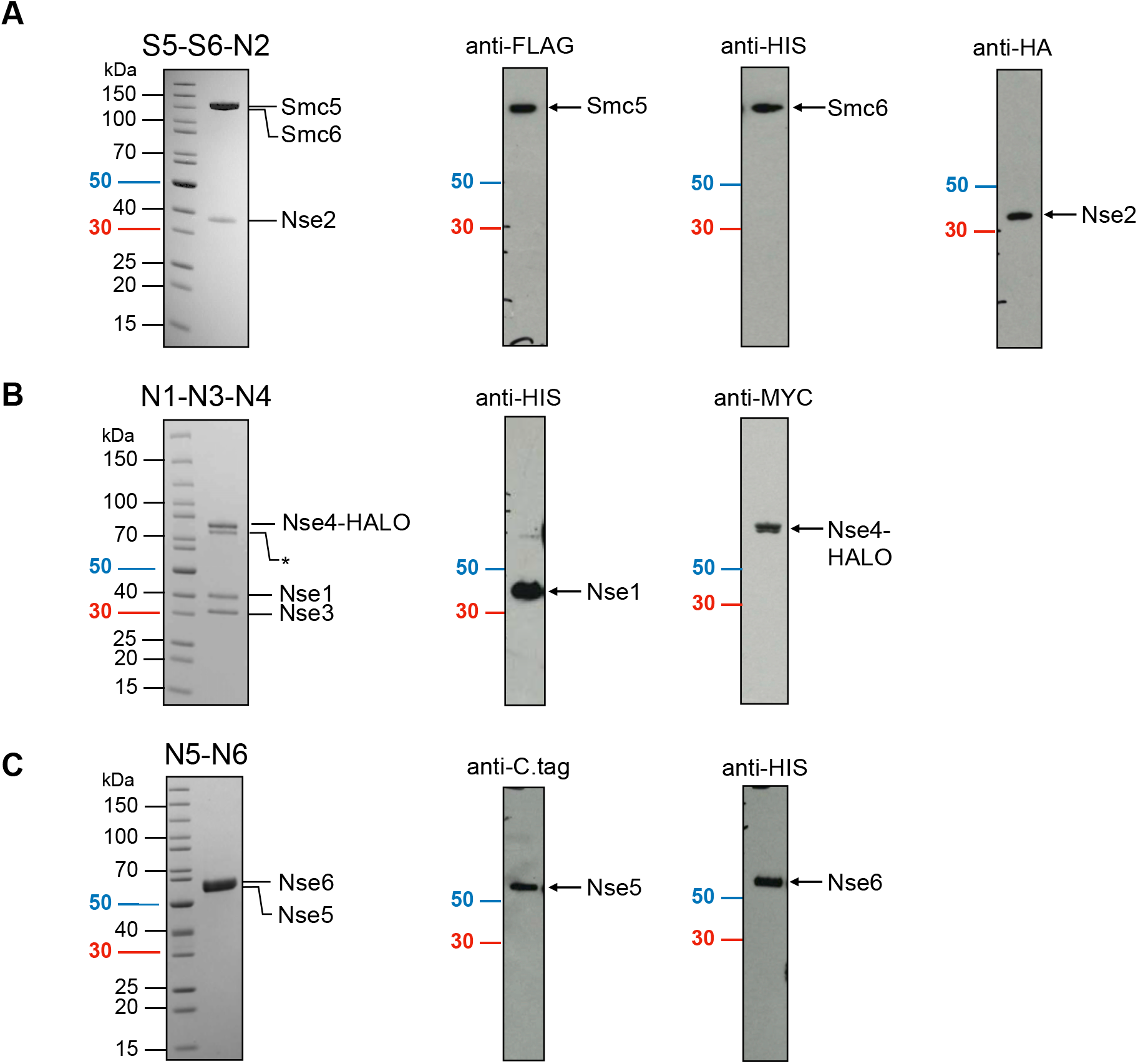
Confirmation of protein identity and migration position on SDS/PAGE gels by western blot. Representative colloidal-blue stained SDS-PAGE gel (left) and associated western blots (right) for each of the purified sub-complexes, expressed by the indicated recombinant baculovirus. (**A**) S5-S6-N2. (**B**) N1-N3-N4. (**C**) N5-N6. The epitope recognised by the primary antibody in each western blot is indicated. To aid comparison, the migration position of the 50 (coloured blue) and 30 kDa (red) molecular mass markers is also highlighted.

**Figure S2.**
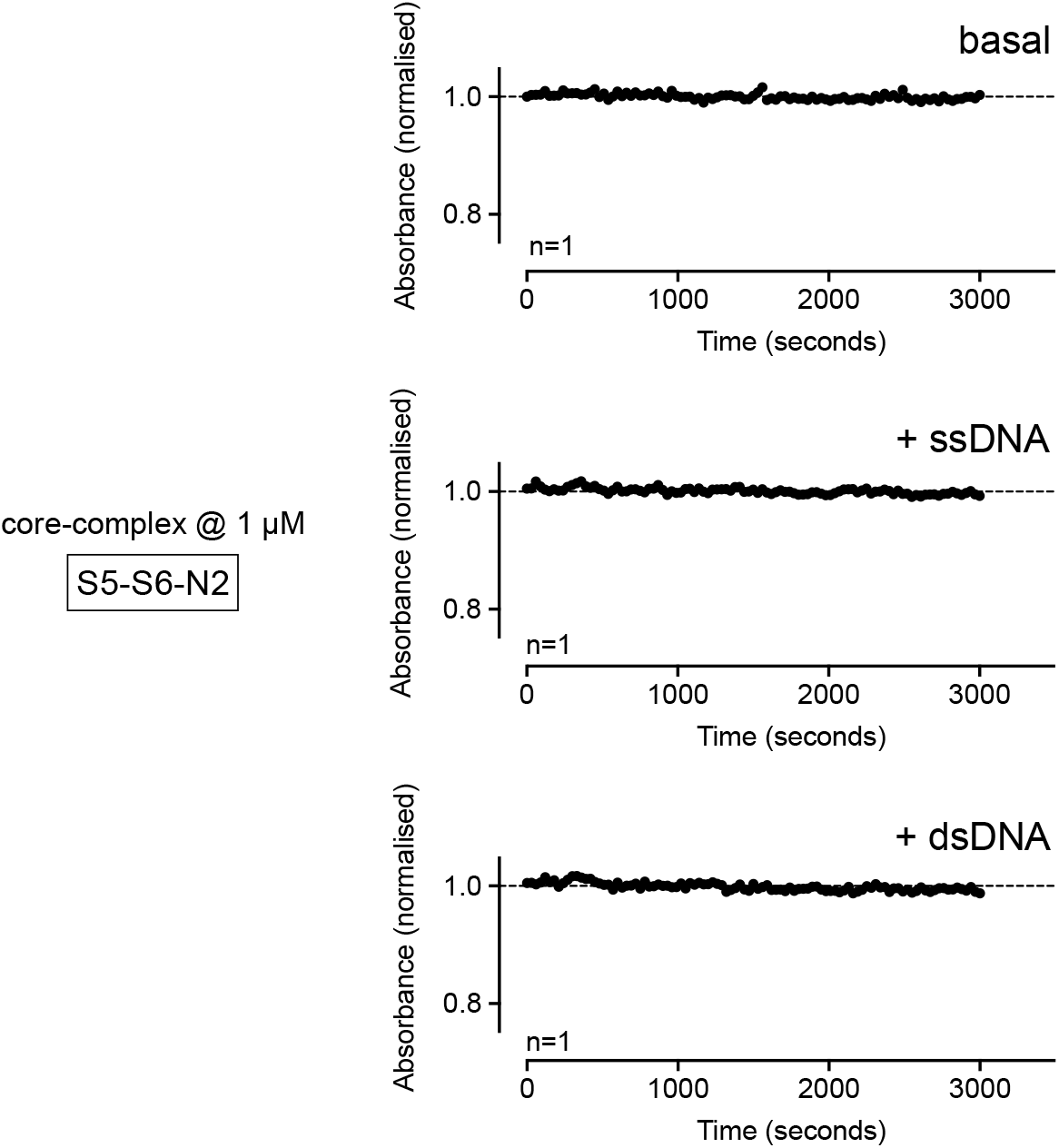
The Smc5/6 ‘core-complex’ does not turnover ATP. Purified Smc5/6 ‘core’ complex, comprising Smc5, Smc6 and Nse2, does not turn over ATP in a NADH-coupled regenerative ATPase assay, even when tested at a final concentration of 1 μM (7-fold higher than the data presented in Figure 5B). Addition of either ssDNA or dsDNA still has no stimulatory effect.

**Figure S3.**
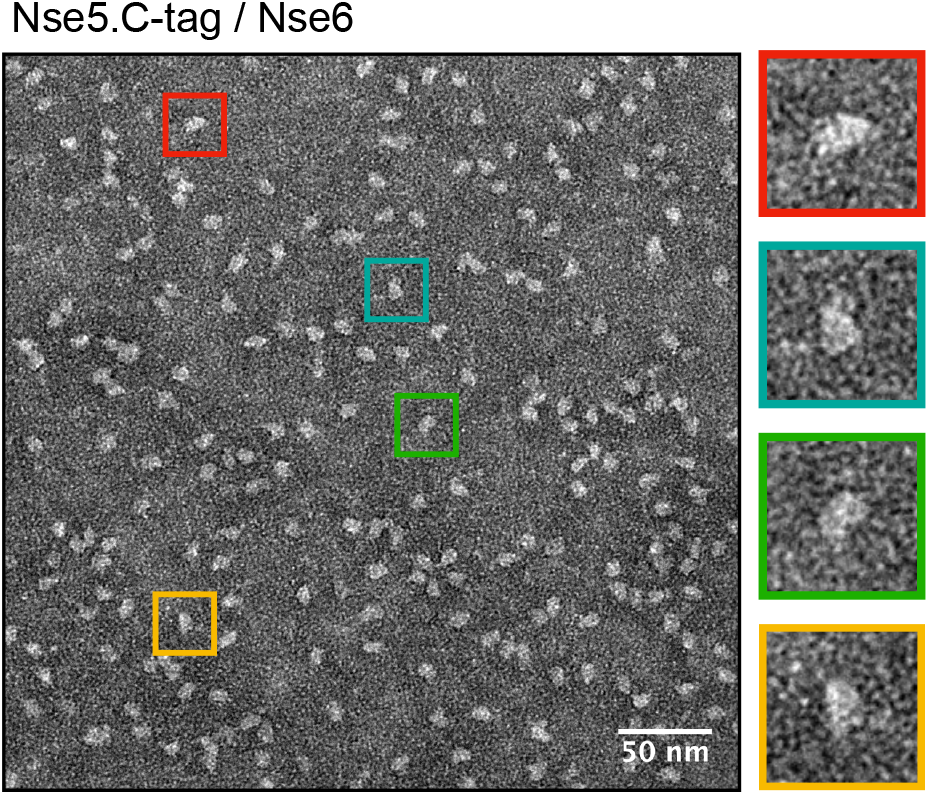
Visualisation of the Nse5/6 heterodimer. Representative micrograph showing particles of purified Nse5/6 heterodimer, negatively stained by uranyl acetate. Selected particles (as indicated by boxes with different coloured borders) are shown at increased magnification on the right-hand side.

**Figure S4.**
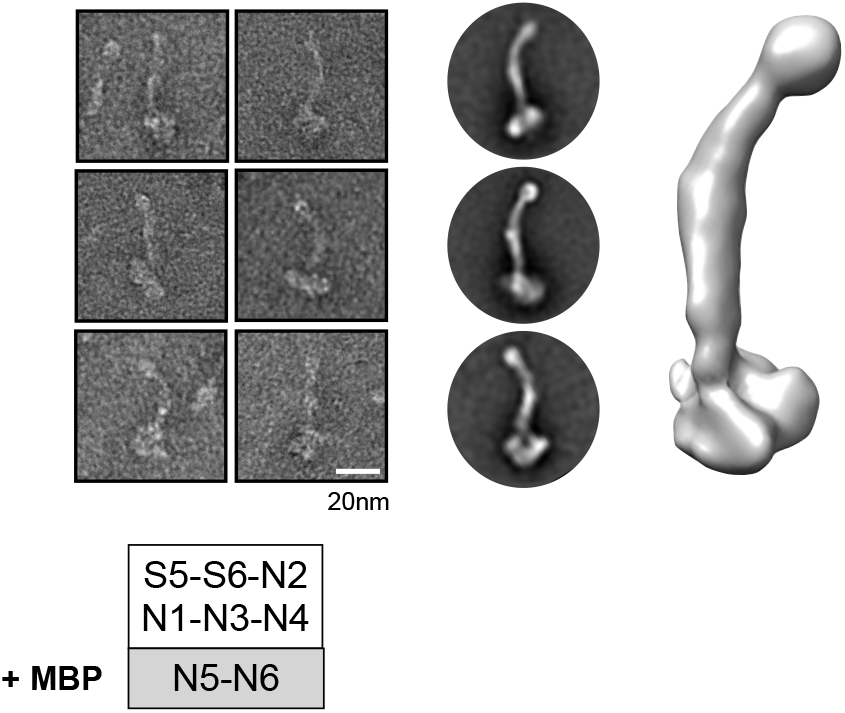
2D class averages and 3D model of the Smc5/6 ‘super-complex’ containing Nse5 fused to MBP. (Left) Representative images of individual particles. (Middle) 2D class averages (Right) Initial 3D reconstruction.

## REFERENCES

1. Alt, A. et al. Specialized interfaces of Smc5/6 control hinge stability and DNA association. Nat Commun 8, 14011 (2017).

2. Griese, J.J., Witte, G. & Hopfner, K.P. Structure and DNA binding activity of the mouse condensin hinge domain highlight common and diverse features of SMC proteins. Nucleic Acids Res 38, 3454–65 (2010).

3. Haering, C.H., Lowe, J., Hochwagen, A. & Nasmyth, K. Molecular architecture of SMC proteins and the yeast cohesin complex. Mol Cell 9, 773–88 (2002).

4. Kurze, A. et al. A positively charged channel within the Smc1/Smc3 hinge required for sister chromatid cohesion. EMBO J 30, 364–78 (2011).

5. Hassler, M., Shaltiel, I.A. & Haering, C.H. Towards a Unified Model of SMC Complex Function. Curr Biol 28, R1266–R1281 (2018).

6. Uhlmann, F. SMC complexes: from DNA to chromosomes. Nat Rev Mol Cell Biol 17, 399–412 (2016).

7. Taylor, E.M., Copsey, A.C., Hudson, J.J., Vidot, S. & Lehmann, A.R. Identification of the proteins, including MAGEG1, that make up the human SMC5-6 protein complex. Mol Cell Biol 28, 1197–206 (2008).

8. Aragon, L. The Smc5/6 Complex: New and Old Functions of the Enigmatic Long-Distance Relative. Annu Rev Genet 52, 89–107 (2018).

9. Diaz, M. & Pecinka, A. Scaffolding for Repair: Understanding Molecular Functions of the SMC5/6 Complex. Genes (Basel) 9(2018).

10. Palecek, J.J. SMC5/6: Multifunctional Player in Replication. Genes 10(2019).

11. Sole-Soler, R. & Torres-Rosell, J. Smc5/6, an atypical SMC complex with two RING-type subunits. Biochemical Society Transactions 48, 2159–2171 (2020).

12. Decorsiere, A. et al. Hepatitis B virus X protein identifies the Smc5/6 complex as a host restriction factor. Nature 531, 386–9 (2016).

13. Murphy, C.M. et al. Hepatitis B Virus X Protein Promotes Degradation of SMC5/6 to Enhance HBV Replication. Cell Rep 16, 2846–2854 (2016).

14. Xu, W. et al. PJA1 Coordinates with the SMC5/6 Complex To Restrict DNA Viruses and Episomal Genes in an Interferon-Independent Manner. J Virol 92(2018).

15. Bentley, P., Tan, M.J.A., McBride, A.A., White, E.A. & Howley, P.M. The SMC5/6 Complex Interacts with the Papillomavirus E2 Protein and Influences Maintenance of Viral Episomal DNA. J Virol 92(2018).

16. Gibson, R.T. & Androphy, E.J. The SMC5/6 Complex Represses the Replicative Program of High-Risk Human Papillomavirus Type 31. Pathogens 9(2020).

17. Payne, F. et al. Hypomorphism in human NSMCE2 linked to primordial dwarfism and insulin resistance. The Journal of clinical investigation 124, 4028–4038 (2014).

18. van der Crabben, S.N. et al. Destabilized SMC5/6 complex leads to chromosome breakage syndrome with severe lung disease. The Journal of clinical investigation 126, 2881–2892 (2016).

19. Hazbun, T.R. et al. Assigning function to yeast proteins by integration of technologies. Mol Cell 12, 1353–65 (2003).

20. Pebernard, S., Wohlschlegel, J., McDonald, W.H., Yates, J.R. 3rd & Boddy, M.N. The Nse5-Nse6 dimer mediates DNA repair roles of the Smc5-Smc6 complex. Mol Cell Biol 26, 1617–30 (2006).

21. Raschle, M. et al. DNA repair. Proteomics reveals dynamic assembly of repair complexes during bypass of DNA cross-links. Science 348, 1253671 (2015).

22. Oravcova, M. & Boddy, M.N. Recruitment, loading, and activation of the Smc5-Smc6 SUMO ligase. Curr Genet 65, 669–676 (2019).

23. Oravcova, M. et al. Brc1 Promotes the Focal Accumulation and SUMO Ligase Activity of Smc5-Smc6 during Replication Stress. Mol Cell Biol 39(2019).

24. Litwin, I., Pilarczyk, E. & Wysocki, R. The Emerging Role of Cohesin in the DNA Damage Response. Genes (Basel) 9(2018).

25. Etheridge, T.J. et al. Single-molecule live cell imaging of the Smc5/6 DNA repair complex. bioRxiv, 2020.06.19.148106 (2020).

26. Bustard, D.E. et al. During replication stress, non-SMC element 5 (NSE5) is required for Smc5/6 protein complex functionality at stalled forks. J Biol Chem 287, 11374–83 (2012).

27. Ohouo, P.Y., Bastos de Oliveira, F.M., Almeida, B.S. & Smolka, M.B. DNA damage signaling recruits the Rtt107-Slx4 scaffolds via Dpb11 to mediate replication stress response. Mol Cell 39, 300–6 (2010).

28. Verkade, H.M., Bugg, S.J., Lindsay, H.D., Carr, A.M. & O’Connell, M.J. Rad18 is required for DNA repair and checkpoint responses in fission yeast. Mol Biol Cell 10, 2905–18 (1999).

29. Sheedy, D.M. et al. Brc1-mediated DNA repair and damage tolerance. Genetics 171, 457–68 (2005).

30. Wan, B., Wu, J., Meng, X., Lei, M. & Zhao, X. Molecular Basis for Control of Diverse Genome Stability Factors by the Multi-BRCT Scaffold Rtt107. Mol Cell 75, 238–251 e5 (2019).

31. Adamus, M. et al. Molecular Insights into the Architecture of the Human SMC5/6 Complex. J Mol Biol 432, 3820–3837 (2020).

32. Duan, X. et al. Architecture of the Smc5/6 Complex of Saccharomyces cerevisiae Reveals a Unique Interaction between the Nse5-6 Subcomplex and the Hinge Regions of Smc5 and Smc6. J Biol Chem 284, 8507–15 (2009).

33. Palecek, J., Vidot, S., Feng, M., Doherty, A.J. & Lehmann, A.R. The Smc5-Smc6 DNA repair complex. bridging of the Smc5-Smc6 heads by the KLEISIN, Nse4, and non-Kleisin subunits. J Biol Chem 281, 36952–9 (2006).

34. Duan, X. et al. Structural and functional insights into the roles of the Mms21 subunit of the Smc5/6 complex. Mol Cell 35, 657–68 (2009).

35. Duan, X. & Ye, H. Purification, crystallization and preliminary X-ray crystallographic studies of the complex between Smc5 and the SUMO E3 ligase Mms21. Acta Crystallogr Sect F Struct Biol Cryst Commun 65, 849–52 (2009).

36. Barnett, J.T. & Kad, N.M. Understanding the coupling between DNA damage detection and UvrA’s ATPase using bulk and single molecule kinetics. FASEB J 33, 763–769 (2019).

37. Elbatsh, A.M.O. et al. Distinct Roles for Condensin’s Two ATPase Sites in Chromosome Condensation. Mol Cell 76, 724–737 e5 (2019).

38. Hassler, M. et al. Structural Basis of an Asymmetric Condensin ATPase Cycle. Mol Cell 74, 1175–1188 e9 (2019).

39. Fousteri, M.I. & Lehmann, A.R. A novel SMC protein complex in Schizosaccharomyces pombe contains the Rad18 DNA repair protein. EMBO J 19, 1691–702 (2000).

40. Kanno, T., Berta, D.G. & Sjogren, C. The Smc5/6 Complex Is an ATP-Dependent Intermolecular DNA Linker. Cell Rep 12, 1471–82 (2015).

41. Kimura, K. & Hirano, T. ATP-dependent positive supercoiling of DNA by 13S condensin: a biochemical implication for chromosome condensation. Cell 90, 625–34 (1997).

42. Murayama, Y. & Uhlmann, F. Biochemical reconstitution of topological DNA binding by the cohesin ring. Nature 505, 367–71 (2014).

43. Anderson, D.E., Losada, A., Erickson, H.P. & Hirano, T. Condensin and cohesin display different arm conformations with characteristic hinge angles. J Cell Biol 156, 419–24 (2002).

44. Burmann, F. et al. A folded conformation of MukBEF and cohesin. Nat Struct Mol Biol 26, 227–236 (2019).

45. Collier, J.E. et al. Transport of DNA within cohesin involves clamping on top of engaged heads by Scc2 and entrapment within the ring by Scc3. Elife 9(2020).

46. Higashi, T.L. et al. A Structure-Based Mechanism for DNA Entry into the Cohesin Ring. Mol Cell 79, 917–933 e9 (2020).

47. Kong, M. et al. Human Condensin I and II Drive Extensive ATP-Dependent Compaction of Nucleosome-Bound DNA. Mol Cell 79, 99–114 e9 (2020).

48. Lee, B.G. et al. Cryo-EM structures of holo condensin reveal a subunit flip-flop mechanism. Nat Struct Mol Biol 27, 743–751 (2020).

49. Shi, Z., Gao, H., Bai, X.C. & Yu, H. Cryo-EM structure of the human cohesin-NIPBL-DNA complex. Science 368, 1454–1459 (2020).

50. Burmann, F. et al. Tuned SMC Arms Drive Chromosomal Loading of Prokaryotic Condensin. Mol Cell 65, 861–872 e9 (2017).

51. Lindroos, H.B. et al. Chromosomal association of the Smc5/6 complex reveals that it functions in differently regulated pathways. Molecular Cell 22, 755–767 (2006).

52. Pebernard, S., Schaffer, L., Campbell, D., Head, S.R. & Boddy, M.N. Localization of Smc5/6 to centromeres and telomeres requires heterochromatin and SUMO, respectively. EMBO J 27, 3011–23 (2008).

53. Minamino, M., Higashi, T.L., Bouchoux, C. & Uhlmann, F. Topological in vitro loading of the budding yeast cohesin ring onto DNA. Life Sci Alliance 1(2018).

54. Petela, N.J. et al. Scc2 Is a Potent Activator of Cohesin’s ATPase that Promotes Loading by Binding Scc1 without Pds5. Mol Cell 70, 1134–1148 e7 (2018).

55. Leung, G.P., Lee, L., Schmidt, T.I., Shirahige, K. & Kobor, M.S. Rtt107 is required for recruitment of the SMC5/6 complex to DNA double strand breaks. J Biol Chem 286, 26250–7 (2011).

56. Li, X. et al. Structure of C-terminal tandem BRCT repeats of Rtt107 protein reveals critical role in interaction with phosphorylated histone H2A during DNA damage repair. J Biol Chem 287, 9137–46 (2012).

57. Williams, J.S. et al. gammaH2A binds Brc1 to maintain genome integrity during S-phase. EMBO J 29, 1136–48 (2010).

58. Zabrady, K. et al. Chromatin association of the SMC5/6 complex is dependent on binding of its NSE3 subunit to DNA. Nucleic Acids Res 44, 1064–79 (2016).

59. Weissmann, F. et al. biGBac enables rapid gene assembly for the expression of large multisubunit protein complexes. Proc Natl Acad Sci U S A 113, E2564–9 (2016).

60. Fairhead, M. & Howarth, M. Site-specific biotinylation of purified proteins using BirA. Methods Mol Biol 1266, 171–84 (2015).

61. Djender, S., Beugnet, A., Schneider, A. & De Marco, A. The Biotechnological Applications of Recombinant Single-Domain Antibodies are Optimized by the C-Terminal Fusion to the EPEA Sequence (C Tag). Antibodies 3(2014).

62. Los, G.V. et al. HaloTag: a novel protein labeling technology for cell imaging and protein analysis. ACS Chem Biol 3, 373–82 (2008).

63. Scheres, S.H. RELION: implementation of a Bayesian approach to cryo-EM structure determination. J Struct Biol 180, 519–30 (2012).

64. Zivanov, J. et al. New tools for automated high-resolution cryo-EM structure determination in RELION-3. Elife 7(2018).

65. Pettersen, E.F. et al. UCSF Chimera--a visualization system for exploratory research and analysis. J Comput Chem 25, 1605–12 (2004).

66. Schrödinger, L. The PyMOL Molecular Graphics System, v2.3.2.

